# Live-cell imaging of circular and long non-coding RNAs associated to FUS pathological aggregates by Pepper fluorescent RNA

**DOI:** 10.1101/2024.06.03.597119

**Authors:** Erika Vitiello, Francesco Castagnetti, Lorenzo Stufera Mecarelli, Eleonora D’Ambra, Paolo Tollis, Giancarlo Ruocco, Pietro Laneve, Elisa Caffarelli, Davide Mariani, Irene Bozzoni

## Abstract

Lately, important advancements in visualizing RNAs in fixed and live cells have been achieved. While mRNA imaging techniques are well-established, the development of effective methods for studying non-coding RNAs (ncRNAs) in living cells are still challenging but necessary, as they cover a variety of function and of intracellular localization, including highly dynamic processes like phase-transition, still poorly studied *in vivo*. Addressing this issue, we tagged two exemplary ncRNAs with the innovative fluorescent RNA (fRNA) Pepper. Specifically, we show circ-HDGFRP3 interaction with p-bodies, we recapitulate its recruitment in pathological FUS aggregates in a dynamic fashion and we super-resolve its distribution in such aggregates via Structured Illumination Microscopy. Moreover, we tracked the long non-coding RNA (lncRNA) nHOTAIRM1, a motor neurons-specific constituent of stress granules (SG), monitoring its behavior throughout the oxidative-stress response in physiological and pathological conditions. Overall, as fRNAs development advances, our work shows a successful use of Pepper for the monitoring of complex processes, as phase-transition, of paradigmatic molecules like circular RNAs (circRNAs) and lncRNAs with super-resolution potential in living mammalian cells.

## INTRODUCTION

Over the years, considerable effort has been spent in developing tools for imaging RNAs. While advances in RNA FISH (Fluorescent *In Situ* Hybridization) have yielded significant results in terms of spatial resolution and multiplexing, a lot of information is missing if it comes to study the RNA dynamics. Approaches based on chemically synthesized probes and genetically encoded probes were developed for targeting endogenous RNAs (Le et al, 2022); however, they encounter limitations determined by the abundance of the target RNA, high background signal and off-target issues (Ma et al., 2017; Tang et al., 2022; Tyagi and Kramer, 1996; Wang et al., 2022; Yang et al., 2019). The most diffused tools for RNA live imaging take advantage of RNA binding proteins that can be fused to a fluorescent molecule, like a fluorescent protein (FP), to target exogenous RNAs of interest (Le et al., 2022). Nonetheless, this approach requires up to 48 FPs tethered to a single RNA molecule, resulting in high hindrance for the target RNA (Le et al., 2022; Peabody, 1993). To overcome these limitations, fluorescent RNAs are improving fast, too. The first developed dye-binding RNA aptamer, Spinach, is able to bind a molecule that mimics the GFP chromophore, remaining non-fluorescent in its free state but becoming fluorescent upon binding to the RNA aptamer, due to rigidification (Paige et al., 2011). In 2019, Yang and colleagues (Chen et al., 2019) described the new family of dye-binding aptamers Peppers, addressing the main issues of previously described fRNAs (Jeng et al., 2016; Paige et al., 2011), which were limited in brightness, colour and photostability. The Pepper aptamer binds a series of custom small molecules named HBC, proving enhanced efficiency in targeting a wide range of RNAs with a relatively short RNA tag, potentially enabling the visualization of different classes of ncRNAs.

Nonetheless, with the exception of circular RNA aptamers utilized as metabolite biosensors (Litke and Jaffrey, 2019), there are no examples in literature of live imaging experiments performed on circRNAs and few examples are provided for lncRNAs (Cawte et al., 2020; Chen et al., 2019; Sarfraz et al., 2023; Yang et al., 2019).

Indeed, circRNAs originate from a non-canonical splicing event, the back-splicing reaction, a process that is driven by the matching of complementary intronic sequences or by specific RNA binding proteins (Ashwal-Fluss et al., 2014; Conn et al., 2015; Errichelli et al., 2017; Fei et al., 2017; Kramer et al., 2015). The back-splicing junction (BSJ) is the only specific sequence that distinguishes a circRNA from other linear isoforms deriving from the same pre-mRNA, making circRNAs challenging to target both in fixed and in live cells (Bejugam et al., 2020).

Interestingly, non-coding RNAs, and in particular circRNAs, are thought to play a major role in ribonucleoparticles (RNPs) assembly and transport and in the promotion of liquid-liquid phase separation (LLPS) events (Corbet and Parker, 2019; D’Ambra et al., 2019; Jain et al., 2016; Khong et al., 2017; Molliex et al., 2015; Namkoong et al., 2018; Rybak-Wolf et al., 2015; You et al., 2015). These biological processes, often de-regulated in neurodegenerative diseases, are highly dynamic (Protter and Parker, 2016), making it crucial to explore them through live imaging assays. While important contributions have already been given in the study of these processes for several mRNA species (Le et al., 2022), the application of live imaging techniques in the context of ncRNAs remains quite limited and unexplored (Yang et al., 2019).

With these premises, we designed overexpression plasmids to track the dynamics of paradigmatic ncRNAs, associated to pathological condensates formed upon mutations in the ALS-linked FUS protein, tagging them with Pepper fluorescent RNA (Chen et al., 2019). Specifically, we dedicated our effort to visualize the circRNA circ-HDGFRP3 (D’Ambra et al., 2021) and the lncRNA nHOTAIRM1 (Rea et al., 2020; Tollis et al., 2023) through live imaging in mammalian cells, providing an effective model for studying diverse classes of ncRNAs in physiological and pathological conditions.

## RESULTS

### Engineering and validation of overexpression constructs for circRNAs live imaging

circ-HDGFRP3, which originates from the circularisation of exons 2, 3, 4 and 5 of the HDGFRP3 (Hepatoma-Derived Growth Factor-Related Protein 3) gene, was selected as a representative example for live imaging due to its ability to traffic along neurites under normal conditions, while being trapped in protein condensates when FUS aggregates are induced (D’Ambra et al, 2021).

The overexpression construct for circ-HDGFRP3 tagged with the fRNA Pepper was obtained by cloning the region to be circularized between the inverted complementary sequences (ICSs) of the ZKSCAN1 MCS vector (Kramer et al., 2015; Legnini et al., 2017), which also harbours a doxycycline inducible promoter (hereby named “p-circ”, Fig 1A). Secondly, to tag the circRNA with Pepper preserving the integrity of the BSJ, we identified two different Pepper insertion sites: one within exon 4 of circ-HDGFRP3, generating the plasmid p-circ-Ex4 and another within exon 5, generating the plasmid p-circ-Ex5. An array of 4xPepper was cloned in each insertion site, optimizing the balance between signal to noise ratio and length of the target.

**Figure 1.**
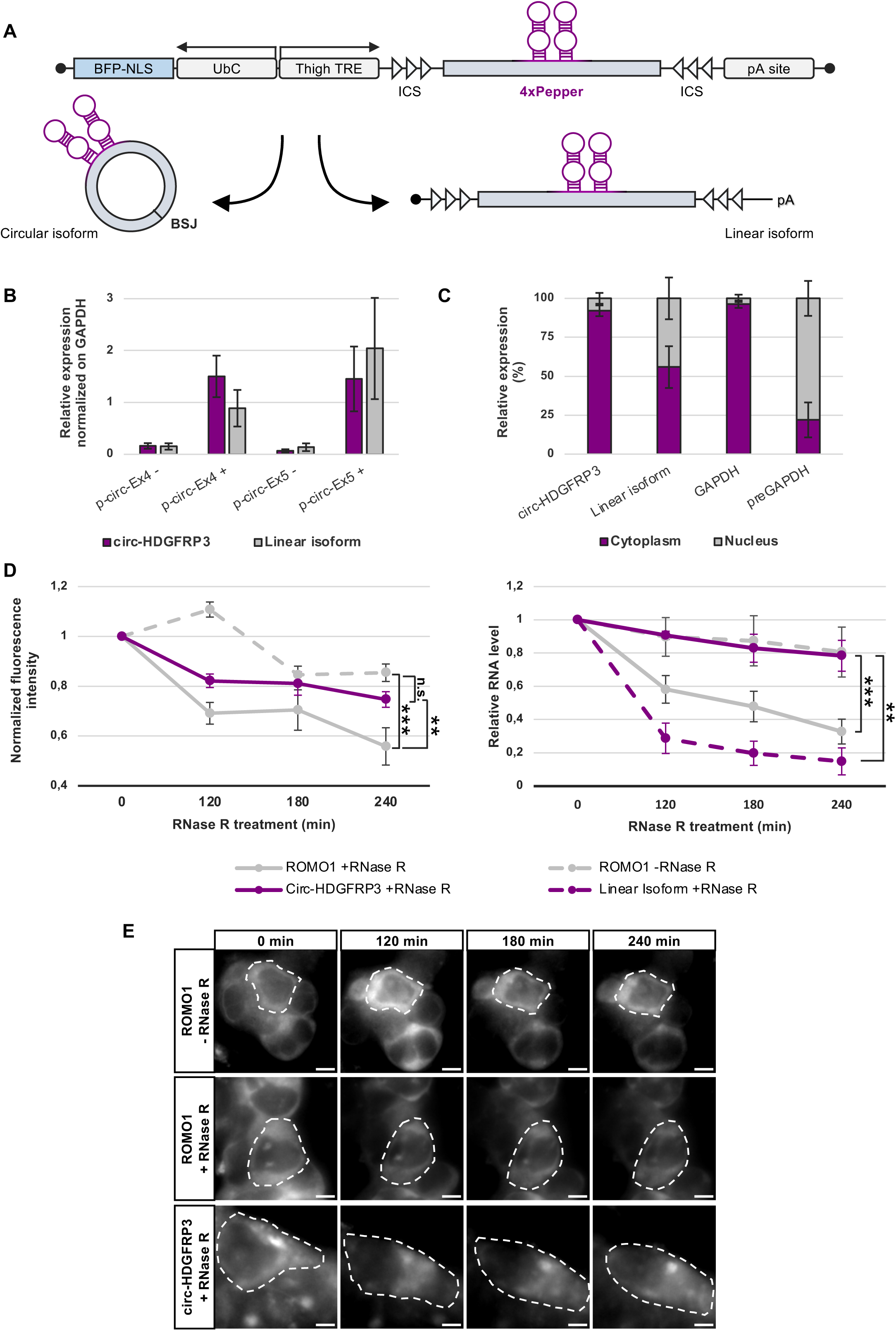
Overexpression constructs for visualization of exogenous circRNAs tagged with Pepper. A Graphical representation of the p-circ overexpression plasmids designed to tag and visualize circRNAs with the fRNA Pepper. B Expression levels quantified through qPCR of circ-HDGFRP3 and of the linear isoform upon p-circ-Ex4 and p-circ-Ex5 transfection, with (+) or without (-) doxycycline administration, normalized on GAPDH levels. Error bars represent ± SEM (N=3). C Nuclear/cytoplasmic fractions of exogenous circ-HDGFRP3 and its relative linear isoform expressed in percentage. GAPDH and preGAPDH were used as cytoplasmic and nuclear controls, respectively. Error bars represent ± SEM (N=3). D Left chart represents normalized fluorescence intensities fluctuations of HBC620 dye bound to ROMO1 mRNA (grey, solid line) or to circ-HDGFRP3 (magenta, solid line) upon 120, 180 and 240 minutes of RNase R treatment. Cells transfected with ROMO1-4xPepper overexpression plasmid but not treated with Triton X-100, hence not exposed to RNase R, were used as a negative control (grey, dashed line). Error bars represent ± SEM. **p = 0.017 and ***p = 0.0000018 correspond to two tails unpaired Student’s t-test (N=3). Right chart represents RNA levels of ROMO1 mRNA exposed (grey, solid line) and not exposed (grey, dashed line) to 120, 180 and 240 minutes of RNase R treatment and levels of circ-HDGFRP3 (magenta, solid line) and of the linear isoform (magenta, dashed line) upon 120, 180 and 240 minutes of RNase R treatment. Error bars represent ± SEM. **p = 0,029 and ***p = 0,000039 correspond to two tails unpaired Student’s t-test (N=3).

After engineering, we tested the overexpression level and the circularization efficiency of the different p-circ constructs. Indeed, the coupling of the ICSs is not a flawlessly efficient process (Kramer et al., 2015) and plasmid transcription could result in a different proportion of properly circularized transcript compared to a spurious linear non-circularized transcript (linear isoform) composed of the ICSs, the exons of the circular RNA, the sequence of Pepper and canonical 5’-cap and poly-A tail (Fig 1A).

Importantly, the linear isoform might as well retain the ability to bind the fluorogenic dye, possibly causing unwanted fluorescent signal.

To determine the expression level of both circular and linear isoforms, qPCR analysis was performed upon transfection and doxycycline induction in HEK 293T cells. qPCR results showed that both p-circ-Ex4 and p-circ-Ex5 ensured satisfactory overexpression of the circular isoform, even if p-circ-Ex5 generated a higher amount of linear isoform compared to p-circ-Ex4. Importantly, p-circ-Ex4 provided a favourable circular-to-linear ratio, with the circular isoform being more abundant than the linear one (Fig 1B).

Therefore, p-circ-Ex4 was selected to evaluate whether the insertion of Pepper affected the localization of exogenous circ-HDGFRP3. Nuclear/cytoplasmic fractionation experiments in HEK 293T cells transfected with p-circ-Ex4 revealed that exogenous circ-HDGFRP3 was mostly exported in the cytoplasm (91%), consistent with the localization of the endogenous transcript, previously described as mainly cytoplasmic (Errichelli et al., 2017). On the other hand, the linear isoform was equally distributed between nucleus (45%) and cytoplasm (55%) (Fig 1C). Altogether, these experiments indicated that Pepper insertion did not perturb the cytoplasmic localization of circ-HDGFRP3.

### Detection of exogenous circ-HDGFRP3 in living cells

To assess the visualization of exogenous circ-HDGFRP3, we treated p-circ-Ex4 transfected HEK 293T cells with the HBC620 fluorogenic dye. Successfully transfected cells were monitored thanks to the tagBFP reporter gene adapted with a nuclear localization signal (BFP-NLS) cloned into the p-circ-Ex4 construct (Fig 1A). To set the background signal, cells without doxycycline were used (Supplementary Fig 1A, left panel). When looking at the HBC620 fluorescent signal in doxycycline-treated cells, we could observe quite clear cytoplasmic focal foci (Supplementary Fig 1A, right panel, yellow arrows), in agreement with the localization of the endogenous circ-Hdgfrp3 (D’Ambra et al., 2021).

To demonstrate that the fluorescent signal corresponding to such foci mainly derived from circularized circ-HDGFRP3 molecules, we treated transfected cells with RNase R and imaged them with HBC620. In fact, it is well described that while circRNAs are resistant to RNase R activity, linear RNAs are rapidly degraded (Nielsen et al., 2022). The transcript of ROMO1 (Chung et al., 2006) tagged with 4xPepper, was used as linear control and indicated that the RNase R treatment was working properly; in fact, while without RNase R we observed only a 15% decrease of the fluorescent signal between 0 and 240 min (Fig 1D, left panel, and 1E), paralleling the reduction in RNA level, which was of 20% (Fig 1D, right panel), upon the nuclease treatment the reduction of the fluorescent signal and of the RNA was quite conspicuous, reaching residual values of 55% and 32% respectively (Fig 1D and 1E). In comparison, the fluorescent signal provided by p-circ-Ex4 after 240 minutes of RNase R treatment decreased only of 25% compared to time 0 (Fig 1D, left panel, and 1E); notably, at the end of the treatment, while the circRNA levels remained at 78% of the initial amount, the linear isoform dropped down to 15% (Fig 1D, right panel). The fact that, in the case of p-circ-Ex4, the fluorescence was only partially affected (25%) upon exonucleolytic treatment, indicated that the detected signals were mostly provided by circular molecules. Moreover, since these cytoplasmic fluorescent signals concentrated in focal foci (Fig 1E), we could derive that these foci were made by circ-HDGFRP3 molecules. Further supporting this hypothesis, we did not identify any discrete puncta in the nuclear compartment, which contains only the linear isoform, allowing us to conclude that such isoform produces diffused signals difficult to discriminate from the background (Fig 1E), which mostly consists of free, unbound HBC (Supplementary Fig 1A, left panel).

### circ-Hdgfrp3 interacts with processing bodies (p-bodies)

To evaluate the identity of such spots, we investigated circ-Hdgfrp3 interaction with canonical cytoplasmic organelles. Since p-bodies are known to act on post-transcriptional regulation of RNAs and to dynamically exchange with stress granules (SG, Moon et al, 2019; Kedersha et al, 2005; Luo et al, 2018) we questioned whether circ-Hdgfrp3 co-localizes with these organelles.

We initially took advantage of Basescope^TM^ technology to perform smFISH on endogenous circ-Hdgfrp3, coupled with IF for the DCP1A (decapping mRNA 1A) protein, a marker of p-bodies and with TIAR protein, a marker of SG. Staining was performed in mESCs (mouse Embryonic Stem Cells) derived motor neurons (MNs) non-treated (Fig 2A, upper panels) or treated with 0.5 mM NaAsO_2_ for one hour, to induce oxidative stress response (Fig 2A, lower panels). A strong association (55%) was observed between circ-Hdgfrp3 and DCP1A signals in non-stressed conditions, suggesting a close interplay between the circRNA and p-bodies (Fig 2B, left chart), while TIAR remained mostly retained in the nucleus (Fig 2A, upper panels). When oxidative stress was induced, 60% of the circ-Hdgfrp3 spots associated with DCP1A (Fig 2B, right chart). In these conditions, we also observed circ-Hdgfrp3 spots localizing with both DCP1A and TIAR (Fig 2A, lower panels, magnified details). Subsequently, we generated HEK 293T stable cell lines expressing DCP1A tagged with eGFP and we verified that exogenous GFP-DCP1A co-localized with endogenous p-bodies, stained with anti-DCP1A antibody (Supplementary Fig 2A) in unstressed (89%) and stressed (83%) conditions (Supplementary Fig 2B). We transfected these cells with the p-circ-Ex4 plasmid and imaged circ-HDGFRP3 and GFP-DCP1A for 15 minutes at a temporal resolution of ∼11sec/frame; then we performed single particle tracking (SPT) on selected Regions Of Interest (ROIs) taking advantage of the TrackMate plug-in on Fiji Image-J (Ershov et al., 2022; Tinevez et al., 2017). Trajectories of representative particles were plotted as a function of time, confirming a persistent association between the circRNA and p-bodies (Fig 2C and D, Supplementary Movie 1). Besides, we also imaged circ-HDGFRP3 and GFP-DCP1A upon NaAsO_2_ treatment and we measured the Pearson’s Correlation Coefficient (PCC) over time. In line with the analysis in fixed samples, since the PCC value fluctuated between 0.2 - 0.3 over time, we could define that circ-HDGFRP3 positively and steadily correlated with GFP-DCP1A (Fig 2E and Supplementary Movie 2).

**Figure 2.**
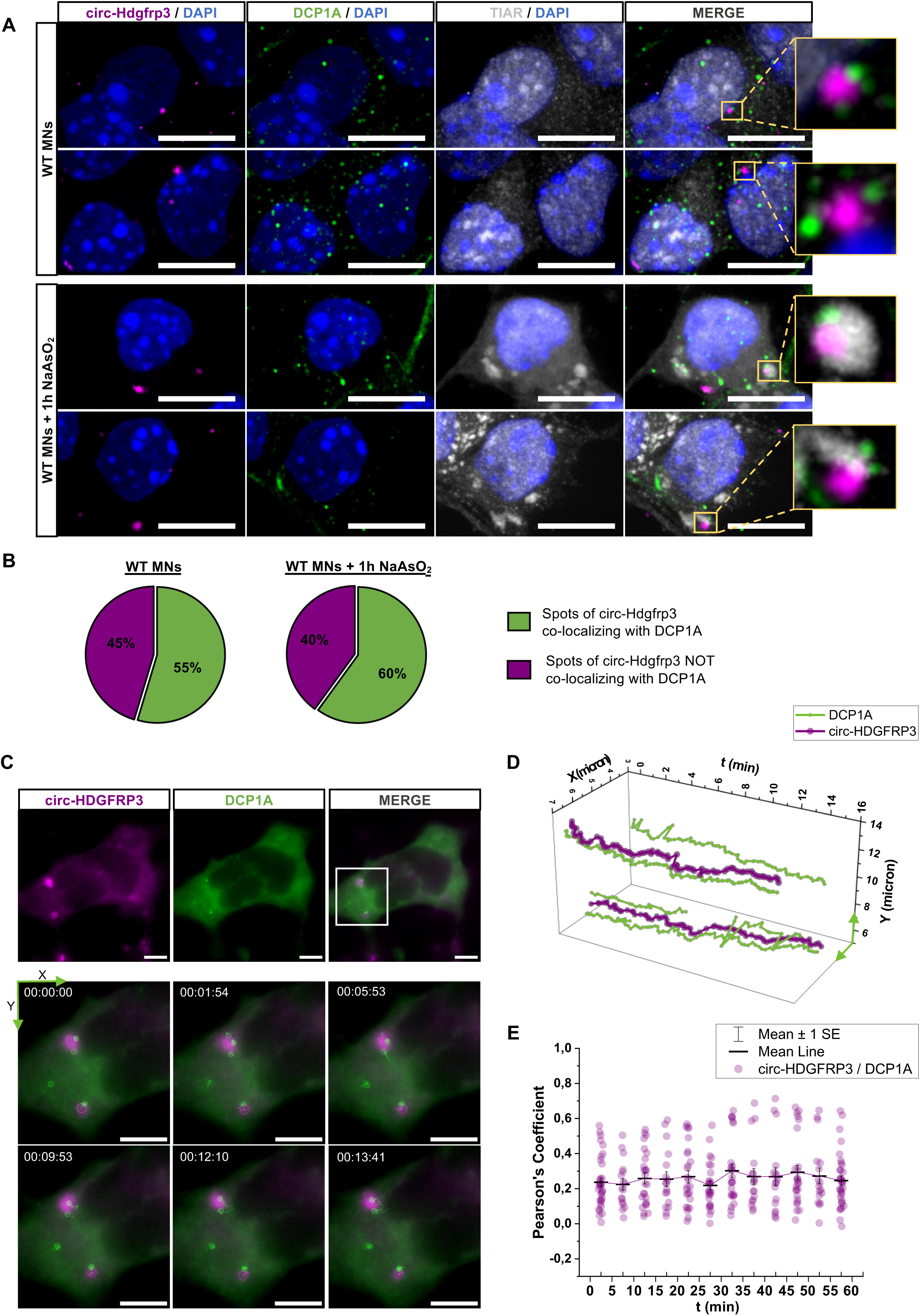
circ-Hdgfrp3 interacts with p-bodies in murine MNs and in living mammalian cells. A Upper panels: two examples of co-localization between circ-Hdgfrp3 (magenta) and DCP1A (green) in non-stressed mESCs-derived MNs. Lower panels: two examples of association between circ-Hdgfrp3 (magenta), DCP1A (green) and TIAR (grey) in stressed mESCs-derived MNs. Oxidative stress response was monitored with TIAR (grey) staining. Magnified yellow boxes highlight the co-localizing spots. Scale bars represent 10 μm. B Quantification of circ-Hdgfrp3 spots co-localizing with DCP1A signals in murine MNs in non-stressed (left panel) and stressed (right panel) conditions. (N_WT MNs_ = 25 cells; N_WT MNs + NaAsO2_= 30 cells). C Representative HEK 293T cells expressing circ-HDGFRP3 labelled with HBC620 (magenta) and GFP-DCP1A (green). Lower panel shows a magnification view of the white box with circ-HDGFRP3 and DCP1A interacting over time (duration 15 min; interval ∼11sec). Spots and tracks detected with TrackMate Plug-in on Fiji-ImageJ are overlaid respectively as magenta (circ-HDGFRP3) and green (DCP1A) circles and lines. All scale bars correspond to 5 μm. D Single particle tracking of circ-HDGFRP3 and DCP1A spots shown in panel C over time. The plot indicates circ-HDGFRP3 (magenta) and DCP1A (green) xy-coordinates (micron) as a function of time (min). E Scatter interval plot representing Pearson’s Correlation Coefficient between circ-HDGFRP3 and DCP1A over time upon NaAsO_2_ administration. Dots represent individual measurements of Pearson’s Correlation Coefficient in an interval of 5 minutes collected from 17 time-lapse acquisitions (duration 1 hour; interval 1-5 min) from 2 independent biological replicates. Horizontal lines indicate mean values and error bars represent ± SEM.

### circ-HDGFRP3 tagged with Pepper fRNA associates to mutant FUS aggregates in living mammalian cells

Next, we wanted to test through live imaging the association of Pepper-tagged circ-HDGFRP3 in FUS aggregates. Hence, we produced stable HEK 293T cell lines expressing FUS^P525L^ tagged with eGFP. Remarkably, when transfecting cells expressing GFP-FUS^P525L^ with p-circ-Ex4, a strong co-localization between the GFP and the HBC620 signal was observed, suggesting that exogenous circ-HDGFRP3 correctly co-localizes with FUS^P525L^ aggregates (Fig 3A and Supplementary Movie 3). We then imaged the particles for ∼10 minutes at a temporal resolution of ∼11sec/frame and we measured the average number of interactions in an interval of one minute (Fig 3B, magenta), showing that circ-HDGFRP3 and FUS^P525L^ signals strongly intersect in both space and time. Moreover, we also performed SPT for both circ-HDGFRP3 and FUS^P525L^ signals, confirming a persistent co-localization between FUS^P525L^ and the circRNA (Fig 3C and Supplementary Movie 3).

**Figure 3.**
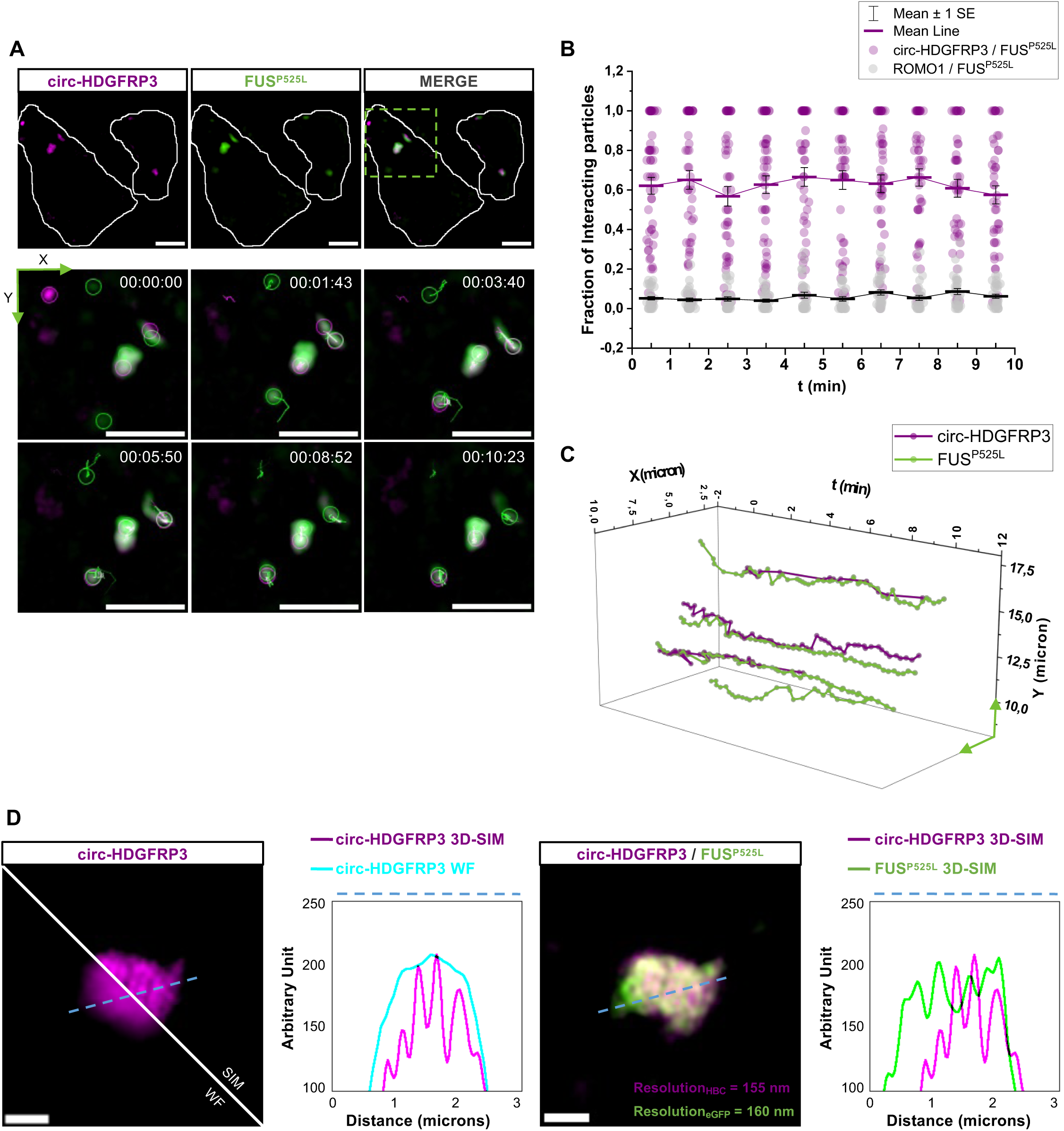
Visualization of exogenous circ-HDGFRP3 co-localizing with FUS^P525L^ aggregates in living HEK 293T cells. A Representative HEK 293T cells transfected with p-circ-Ex4 expressing both circ-HDGFRP3 labelled with HBC620 (magenta) and GFP-FUS^P525L^ (green). Lower panel shows a magnification view of the green dotted box with some circ-HDGFRP3 and FUS^P525L^ particles interacting over time (duration 10 min; interval ∼11sec). Spots and tracks detected with TrackMate Plug-in on Fiji-ImageJ are overlaid respectively as magenta (circ-HDGFRP3) and green (FUS^P525L^) circles and lines. All scale bars correspond to 5 μm. B Scatter interval plot representing circ-HDGFRP3 (magenta) and ROMO1 (grey) fraction of interacting particles with FUS^P525L^ over time. Dots represent individual measurements of number of interacting particles/total number of particles per cell in an interval of 1 min. Error bars represent ± SEM. Magenta and black lines represent mean values. Average and SEM values were calculated from 10 time-lapse measurements (duration 10 min; interval 11-13 sec) collected from 4 independent biological replicates for circ-HDGFRP3/FUS^P525L^ interactions and from 6 measurements from 2 independent biological replicates for ROMO1/FUS^P525L^ interactions. C Single particle tracking of circ-HDGFRP3 and FUS^P525L^ particles shown in panel A over time. The plot indicates circ-HDGFRP3 (magenta) and FUS^P525L^ (green) xy-coordinates (micron) as a function of time (min). D Representative FUS^P525L^ aggregate containing circ-HDGFRP3. Starting from left, signals of circ-HDGFRP3 labelled with HBC620 imaged with Widefield or with 3D-SIM are compared. Intensity profiles (Arbitrary Unit) along the reference blue dotted line of circ-HDGFRP3 signal imaged with Widefield (cyan) or with 3D-SIM (magenta) microscopy are plotted in second panel. Third panel shows FUS^P525L^ signal (green) merged with circ-HDGFRP3 signal (magenta) imaged with 3D-SIM. Respective resolution values calculated on the image are indicated at the bottom of the image. In the last panel intensity profiles (Arbitrary Unit) of circ-HDGFRP3 and FUS^P525L^ imaged with 3D-SIM along the blue dotted line are represented. All scale bars correspond to 1 μm.

To assess that the observed association was actually due to the circRNA sequence and was not caused by a spurious interaction between the sequence of Pepper and the FUS^P525L^ protein, we also performed live imaging and SPT of ROMO1 tagged with four Pepper repetitions. Indeed, this mRNA was selected as a negative control since it is strongly depleted from both WT and FUS^P525L^-containing SG in the dataset of Mariani et al, 2023. The average number of interactions in a one-minute interval was counted (Fig 3B, grey) and trajectories of ROMO1 and FUS^P525L^ were tracked and plotted (Supplementary Fig 3A and B, Supplementary Movie 4). These measurements showed a very low interaction between ROMO1 mRNA and FUS^P525L^, confirming that Pepper tag alone is not sufficient to determine a stable interaction with FUS^P525L^.

Moreover, since the HBC620 dye is compatible with structured illumination microscopy (SIM), we super-resolved the localization of circ-HDGFRP3 in the larger FUS^P525L^ aggregates. As shown by the intensity profiles, a substantial increase in axial resolution was obtained imaging Pepper-tagged circ-HDGFRP3 by SIM, when compared to Widefield microscopy (Fig 3D, left panels). Moreover, by plotting the intensity profile of both HBC620 and eGFP signals imaged with SIM, we were able to determine circ-HDGFRP3 sub-compartmental distribution in FUS^P525L^ aggregates, unveiling non-uniform signal foci of the circRNA partaking in such structures (Fig 3D, right panels). Unequal distribution of RNA in phase-separated granules imaged in super-resolution has already been described in other systems (Jain et al., 2016). Noticeably, the foci of circ-HDGFRP3 signals did not show perfect co-planarity with the ones of FUS^P525L^, in line with the assumption that, as predicted by CatRapid (Bellucci et al, 2011), there are no binding sites for FUS on circ-HDGFRP3. Therefore, these measurements suggest that circ-HDGFRP3 entrapment in pathological FUS aggregates might be due to the intermediation of different protein partners.

Taken together, these data demonstrate that Pepper is a potent tool for live, super-resolved imaging of circRNAs, when associated to specific membrane-less organelles, enabling the monitoring of their intracellular localization and interaction in supra-molecular structures.

### The lncRNA nHOTAIRM1 localizes in physiological and pathological SG

Many lncRNAs have been linked to the formation of amyloid aggregates and to the pathogenesis of diverse neurodegenerative diseases (Wang et al., 2018). For this reason, we also decided to explore the behaviour of nHOTAIRM1, an attractive case study of lncRNA in an ALS-like context. The neuronal isoform of the lncRNA HOTAIRM1 (nHOTAIRM1) plays a crucial role in the control of neuronal differentiation (Rea et al., 2020; Tollis et al., 2023) and is highly expressed in MNs. nHOTAIRM1 has been shown to directly interact with the FUS protein in the cytoplasm (Rea et al., 2020), where FUS regulates its abundance. Nonetheless, it remained un-investigated how nHOTAIRM1 responds to stress stimuli, specifically in ALS conditions.

Therefore, we examined its behaviour in WT and FUS^P525L^ human iPSCs-derived MNs upon NaAsO_2_ administration (De Santis et al., 2018). To do that, we combined smFISH and immunofluorescence, targeting nHOTAIRM1, the SG marker G3BP1 and FUS. As previously reported, we observed that nHOTAIRM1 signal was distributed both in the nucleus and in the cytoplasm (Rea et al., 2020; Tollis et al., 2023), in both WT and FUS^P525L^ conditions (Fig 4A). Co-localization analysis on both nHOTAIRM1 and G3BP1 signals showed that, in stressed WT MNs, a substantial portion of nHOTAIRM1 (64%) co-localized with G3BP1-positive SG (Fig 4B), while, as expected, WT FUS mostly localizes in the nucleus (Fig 4A, upper panel) (Dormann et al., 2010; Kino et al., 2011). In FUS^P525L^ MNs, where FUS de-localizes in the cytoplasm and participates in SG (Fig 4A, lower panel) (Bosco et al., 2010; Lenzi et al., 2015), nHOTAIRM1 was again recruited to SG. Specifically, 45% of the spots co-localized with SG positive for both G3BP1 and mutant FUS, 19% of the signal co-localized with G3BP1 alone, 4% co-localized only with mutant FUS and 32% of the spots did not co-localize (Fig 4C). Collectively, these data indicate that nHOTAIRM1 is recruited to SG upon oxidative stress induction independently of FUS mutation. Indeed, both WT and FUS^P525L^ MNs exhibited the same percentage of nHOTAIRM1 localized in SG, with no significant difference between the two conditions (Fig 4B and C).

**Figure 4.**
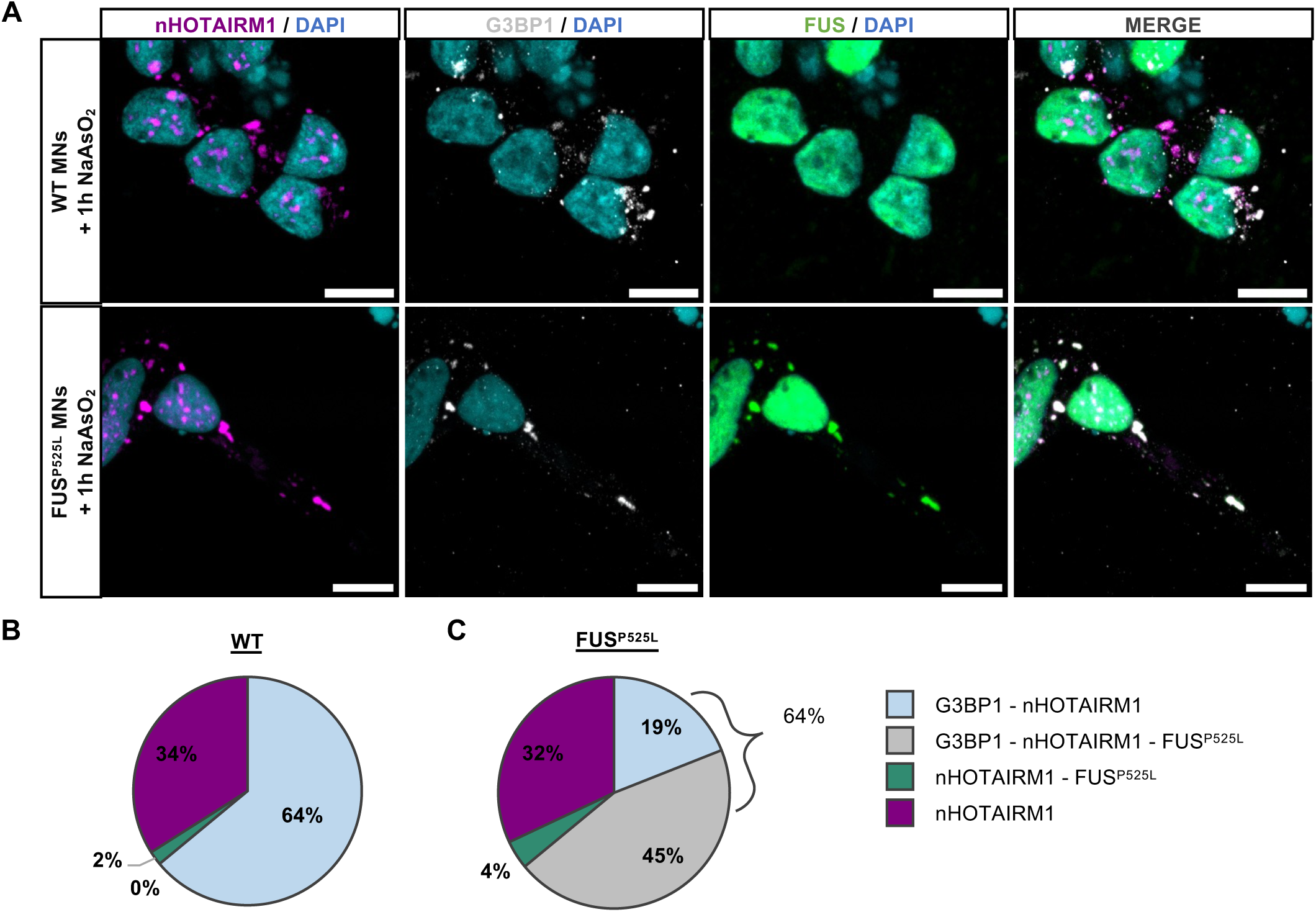
The lncRNA nHOTAIRM1 co-localizes with physiological and pathological SG. A Combination of smFISH and IF for the detection of nHOTAIRM1 (magenta), G3BP1 (grey) and FUS (green) in stressed WT (upper panel) or stressed FUS^P525L^ (lower panel) iPSCs derived MNs. Nuclei stained with DAPI are shown in cyan. Scale bars correspond to 10 μm. B, C Pie charts representing nHOTAIRM1 percentage of co-localization in WT (B) or FUS^P252L^ (C) MNs with G3BP1-positive granules (light blue), FUS-positive granules (green), G3BP1- and FUS^P525L^-positive granules (grey) and percentage of free nHOTAIRM1 spots (magenta). Mean percentages from 3 independent biological replicates are shown (N_WT_ = 549 cells; N ^P525L^ = 84 cells).

Moreover, we stained WT and FUS^P525L^ MNs, treated or non-treated with NaAsO_2_, with TIAR and DCP1A, together with the smFISH for nHOTAIRM1 (Supplementary Fig 4A) and we calculated PCC between nHOTAIRM1 and DCP1A signal to determine if nHOTAIRM1 interacts with p-bodies. A PCC close to 0 was measured in unperturbed WT MNs (Supplementary Fig 4B), indicating a poor correlation between nHOTAIRM1 and p-bodies. The PCC increased significantly when inducing oxidative stress (Supplementary Fig 4B), in line with the assumption that nHOTAIRM1 co-localizes with SG (Fig 4B and Supplementary Fig 4C) and that p-bodies dock onto SG upon NaAsO_2_ treatment (Supplementary Fig 4D) (Kedersha et al., 2005). Interestingly, the PCC between nHOTAIRM1 and DCP1A increased in FUS^P525L^ conditions regardless of stress induction (Supplementary Fig 4B). As it was previously demonstrated that nHOTAIRM1 biochemically interacts with FUS (Rea et al., 2020; Tollis et al., 2023) and that mutant FUS binds and sequesters proteins involved in p-bodies formation (Takanashi and Yamaguchi, 2014), these data suggest that the presence of mutant FUS in the cytoplasm could drive the interaction between nHOTAIRM1 and p-bodies.

### nHOTAIRM1 is dynamically recruited in SG in live mammalian cells

Evidence of nHOTAIRM1 recruitment into SG collected in fixed samples was complemented with temporal and dynamic information, leveraging Pepper once again. We designed an overexpression construct to tag nHOTAIRM1 with a 4xPepper array inserted at the 3’ end of the lncRNA (Supplementary Fig 5A). To follow nHOTAIRM1 behaviour during the oxidative stress response, we transfected with the construct in HEK 293T cells stably expressing G3BP1 or FUS^P525L^ tagged with eGFP. The nHOTAIRM1-4xPepper plasmid provided high transcript levels in both HEK 293T stable lines if compared to the housekeeping mRNA for GAPDH (Supplementary Fig 5B). Hence, upon transfection, it was possible to visualize nHOTAIRM1 bound to the HBC620 dye, with a considerable proportion of the total transfected cells emitting a robust fluorescent signal, mostly diffused in the cytoplasm (Fig 5A and D, magenta). Transfecting nHOTAIRM1 tagged with Pepper in GFP-G3BP1 HEK cells, allowed us to observe its dynamics throughout the entire stress event. As expected, within the first 30 minutes upon NaAsO_2_ treatment, we witnessed LLPS of G3BP1, which formed droplet-like structures eventually becoming SG (Fig 5A and B, Supplementary Movie 5). Interestingly, as highlighted in the magnified detail (Fig 5B), nHOTAIRM1 underwent condensation and we could observe fusion events between the lncRNA and the protein concurrent with the formation of mature SG (Fig 5B, yellow arrows). These observations were consistent with an increase over time of Manders’ overlap Coefficient between nHOTAIRM1 and G3BP1 signals (Fig 5C). As Manders’ coefficient can range from 0 to 1, these measurements indicate that a substantial fraction of nHOTAIRM1 signal overlaps with G3BP1 one, upon NaAsO_2_ treatment. As a negative control, we exploited again the mRNA of ROMO1 tagged with 4xPepper and monitored its behaviour upon oxidative stress (Supplementary Fig 5C). In line with the assumption that ROMO1 mRNA is excluded from SG, the average Manders’ coefficient between ROMO1 and G3BP1 remained close to 0 and, importantly, it did not increase over time (Supplementary Fig 5E), confirming minimal pixel co-occurrence between ROMO1 and G3BP1 throughout the whole stress event.

**Figure 5.**
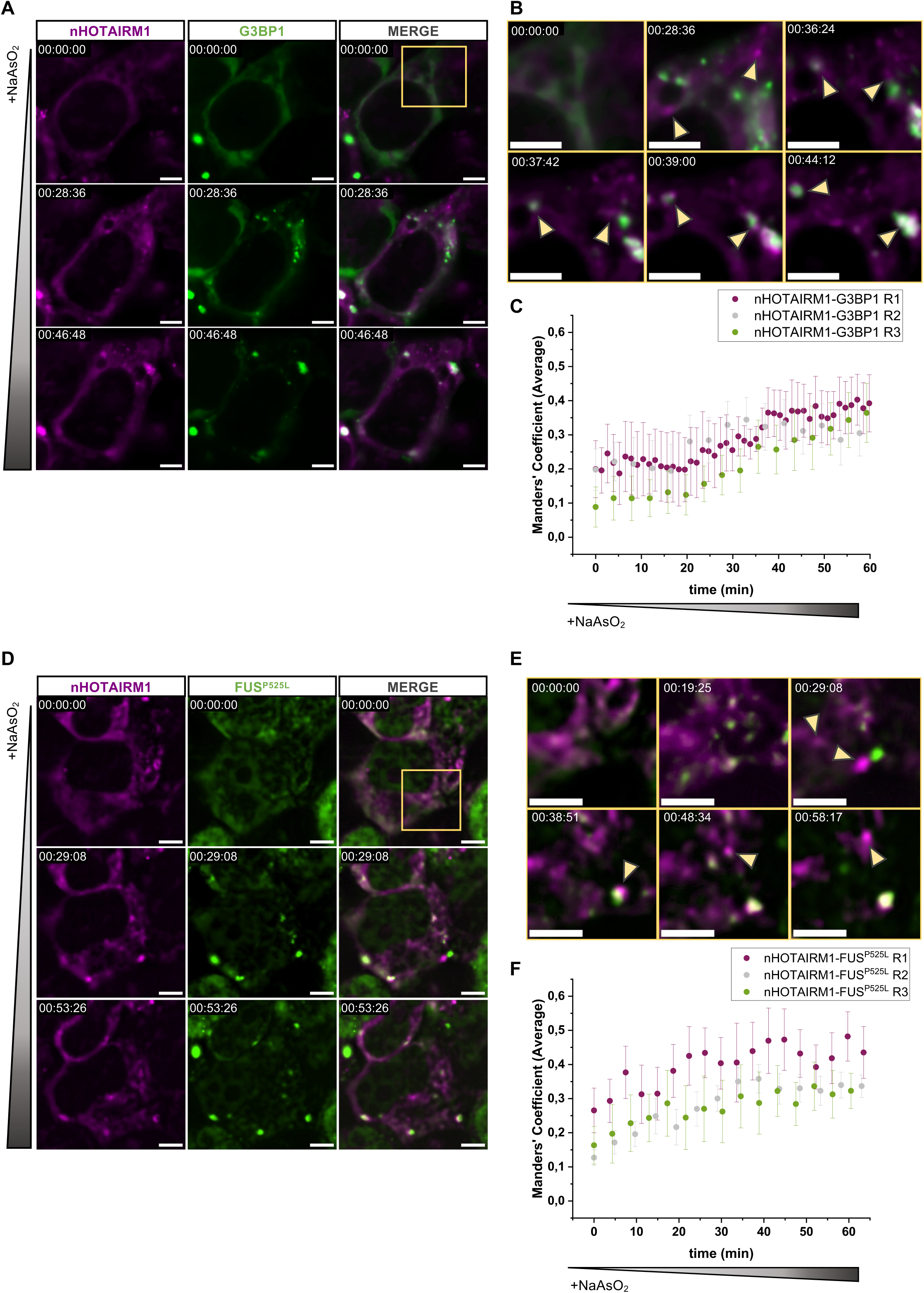
nHOTAIRM1 recruitment in WT or ALS-like granules followed with Pepper fRNA. A Representative cell expressing both nHOTAIRM1 labelled with HBC620 (magenta) and GFP-G3BP1 (green). Time 00:00:00 corresponds to 0.5 mM NaAsO_2_ administration (upper row), while in following time points (second and third row) formation of SG can be followed (duration 1h; interval 1-3 min). All scale bars correspond to 5 μm. B Magnification view of yellow box in panel A showing nHOTAIRM1 and G3BP1 undergoing LLPS throughout the oxidative stress event (yellow arrows) and merging (time points 00:39:00 and 00:44:12). All scale bars correspond to 5 μm. C Average Manders’ Coefficient between nHOTAIRM1 and G3BP1 over time upon NaAsO_2_ administration. Dots represent average values of Manders’ Coefficient. Error bars represent ± SEM. Average and SEM values were calculated from 5 to 10 time-lapse measurements (duration 1 hour; interval 1-5 min) collected from 3 independent biological replicates. Each colour (magenta, grey and green) corresponds to a single biological replicate. D Representative cell expressing both nHOTAIRM1 labelled with HBC620 (magenta) and GFP-FUS^P525L^ (green). Time 00:00:00 corresponds to 0.5 mM NaAsO_2_ administration (upper row), while in following time points (second and third row) formation of FUS^P525L^ condensates can be followed (duration 1h; interval 1-5 min). All scale bars correspond to 5 μm. E Magnification view of yellow box in panel D showing nHOTAIRM1 and FUS^P525L^ first undergoing LLPS throughout the oxidative stress event (yellow arrows) and finally merging (time point 00:38:51). All scale bars correspond to 5 μm. F Average Manders’ Coefficient between nHOTAIRM1 and FUS^P525L^ over time upon NaAsO_2_ administration. Dots represent average values of Manders’ Coefficient. Error bars represent ± SEM. Average and SEM values were calculated from 5 to 10 time-lapse measurements (duration 1 hour; interval 1-5 min) collected from 3 independent biological replicates. Each colour (magenta, grey and green) corresponds to a single biological replicate.

As we determined that nHOTAIRM1 also co-localizes with SG containing FUS^P525L^ (Fig 4C), we replicated the same experiment in HEK 293T cells expressing GFP-FUS^P525L^. Upon stress induction, also FUS^P525L^ underwent condensation in droplet-like structures, mirroring nHOTAIRM1 behaviour. Looking at the interplay between HBC and eGFP signals (Fig 5D and E, Supplementary Movie 6), we observed the same dynamic depicted between nHOTAIRM1 and G3BP1. In parallel, Manders’ coefficient between nHOTAIRM1 and FUS^P525L^ increased over time (Fig 5F), whereas Manders’ coefficient between ROMO1 and FUS^P525L^ remained close to 0 and unaffected by NaAsO_2_ treatment (Supplementary Fig 5D and F).

## DISCUSSION

In summary, we found Pepper (Chen et al., 2019) to be an effective tool to investigate the localization and dynamics of ncRNAs in membrane-less organelles related to phase transition and neurodegenerative-linked processes in living cells. In fact, we successfully reproduced the localization of exogenous circ-HDGFRP3 and nHOTAIRM1 in multiple biomolecular condensates, including SG, p-bodies and pathological FUS^P525L^ aggregates in living cells, recapitulating smFISH experiments on their endogenous counterparts in fixed samples.

In particular, we employed Pepper to follow the dynamics of how the lncRNA nHOTAIRM1 undergoes condensation and fusion events as it is recruited in physiological and pathological granules. Even if the fRNA Pepper proved very straightforward to image linear RNAs, the application to circular RNAs was more challenging. Since circRNAs share their whole sequence with their linear counterparts, except for the back-splicing junction, producing overexpression plasmids for circRNA tagging requires careful design to select the optimal insertion sites and to promote efficient circularization instead of linear counterpart production. Nevertheless, we were able to prove the specificity of circ-HDGFRP3 localization to discrete focal foci through the use of RNase R digestion and we demonstrated its association to p-bodies in physiological conditions as well as its association to FUS aggregates in pathological conditions. Notably, we also explored its sub-compartmental distribution in FUS^P525L^ aggregates through SIM analysis, providing the first instance of a circular RNA being detected with a fluorescent RNA compatible with a super-resolution technique *in vivo* (Bejugam et al., 2020). In fact, to the best of our knowledge, SIM microscopy has not been previously employed for visualizing circRNAs in living samples. Importantly, new generation fRNA such as Pepper and RhoBAST (Chen et al., 2019; Sunbul et al., 2021) are not only compatible with SIM, but also with more sophisticated single-molecule localization microscopy (SMLM) techniques, promising exciting opportunities for the study of the dynamics and the sub-cellular localization of many different RNA species.

## MATERIAL AND METHODS

### Plasmid construction

To produce the p-circ plasmids the described steps were followed: a doxycycline-inducible backbone endowed with flanking inverted complementary sequences (ICSs) (Kramer *et al*, 2015; Legnini *et al*, 2017) for enhanced overexpression of the circular RNAs was already present in the lab. Starting from this plasmid, the In-Fusion HD Cloning kit (Takara Bio) was used to replace the sequence for puromycin resistance with the sequence of the Blue Fluorescent Protein (tagBFP), amplified from the Addgene plasmid (#55312). The SV40 nuclear localization sequence (CACTTTCCGCTTTTTCTTTGG, Addgene plasmid #39319) was cloned downstream the tagBFP combining inverse PCR and blunt end ligation (T4 Ligase NEB). The sequence of circ-HDGFRP3 (exons 2-3-4-5, 522 bp long) was PCR-amplified from SK-N-BE cells cDNA and cloned between the ICSs using In-fusion Cloning Kit (Takara Bio). 4xPepper array sequence (196 bp long) was amplified from the pAPU6-MCS-Pepper (PAPU604MCS1 FR Biotechnology©) plasmid and cloned within exon 4 (between base 43-44) and exon 5 (between base 71-72) of the circ-HDGFRP3 sequence by In-Fusion Cloning Kit (Takara Bio) to obtain the final constructs p-circ-Ex4 and p-circ-Ex5.

The ePB-bsd-Eif1a (PiggyBac transposable vector) plasmid for GFP-G3BP1 expression (Perego et al., 2023) served as a template to generate the constructs for GFP-FUS^P525L^ and GFP-DCP1A expression, using sequential In-Fusion Cloning (Takara Bio) reactions. The same strategy was used to generate epb-bsd-Eif1a plasmids for the overexpression of nHOTAIRM1 and ROMO1 tagged with 4xPepper. CloneAmp HiFi PCR Premix (Clontech) was used for all the PCR amplifications.

### Cell maintenance and manipulation

HEK 293T cells stored at -80°C in freezing medium (DMEM high glucose, Sigma-Aldrich, supplemented with 20% FBS, Gibco, and 10% DMSO), were quickly defrosted at 37°C, using a thermostatic bath. After removing the freezing medium by centrifugation (5 min at 1000 rpm), cells were resuspended in the appropriate amount of maintaining medium (DMEM high glucose, Sigma-

Aldrich, supplemented with 10% FBS, 2 mM GlutaMAX, Gibco, and 1% Penicillin/Streptomycin, Sigma-Aldrich) and plated for maintenance and amplification. For live imaging experiments, the common DMEM high glucose was replaced with FluoroBrite™ DMEM (Thermo Fisher).

To produce HEK 293T cell lines stably expressing GFP-G3BP1, GFP-FUS^P525L^ and GFP-DCP1A, 2.5 × 10^5^ cells were plated on 6 cm dishes. The next day, cells were transfected with a mix of Optimem (Thermo Fisher), 5 μg of specific plasmid, 0.5 μg of hybrid transposase plasmid (Yusa et al., 2011) and Lipofectamine 2000 transfection reagent (Invitrogen™) with a 1:2.5 DNA:transfection reagent ratio.

Cells were then grown in 10 μg/ml blasticidin (Thermofisher Scientific) supplemented medium for 7-10 days to select a successfully transfected pool of cells.

### Cell transfection and induction

For live imaging experiments, 5 × 10^4^ HEK 293T cells per well were plated the day before transfection in 8-well Nunc™ Lab-Tek™ II Chambered Coverglass, previously coated with Geltrex (Thermofisher Scientific) by incubation at 37°C for at least 3 hours. Similarly, for expression and circularization efficiency experiments, 2 × 10^4^ HEK 293T cells were plated two days before transfection on 12-multiwell plate (Corning) coated with 500 μl of Attachment factor X (Gibco) and incubated for 30 min at room temperature. For nucleus-cytoplasm fractionation experiments, 3.5 × 10^5^ cells were instead plated on 6 cm dish (Corning), always two days before transfection.

In both live imaging and circularization efficiency experiments, cells were transfected using a mix of Optimem (Thermo Fisher), 1:2 DNA:FuGENE^®^ HD transfection reagent (Promega) ratio and 2 μg of specific plasmid, while for nucleus-cytoplasm fractionation experiment the amount of transfected plasmid was 2 μg/ml. In case of circular RNAs overexpression experiments, circRNAs transcription was induced using doxycycline at a final concentration of 2 μg/ml, the day after transfection. Doxycycline induction was maintained for 24 hours before cell collection.

### RNA extraction, reverse transcription and qPCR analysis

To evaluate overexpression and circularization efficiency, cells were collected and RNA was extracted using Direct-zol RNA mini-prep kit (Zymo research) following manufacturer instructions 24/48 hours after transfection of overexpression constructs. Residual genomic and plasmidic DNA was removed using DNA-free kit (Invitrogen^TM^) and 500 ng of RNA were retro-transcribed with PrimeScript RT Master Mix (Takara Bio) following manufacturer instructions. cDNA obtained from retro-transcription was analysed by qPCR using the PowerUp SYBR Green Master Mix (Thermo Fischer) reagent coupled with a Quant-Studio 5 (Applied Biosystems) machine. For each reaction 5 ng of cDNA, 7.5 μl of SYBR Green, 0.5 μl of each primer (330 μM as final concentration) and ddH_2_O up to 15 μl of total reaction volume were used. Three technical replicates for each selected target were analysed on a96 well plate (Applied Biosystems™).

For overexpression and circularization efficiency experiments, analysis of qPCR data was conducted as follow: first, the Delta Ct between the target RNA and the reference gene GAPDH was calculated. Those values were then used to calculate the Fold change (FC), to obtain the relative expression of the specific RNA isoform compared to a housekeeping gene. The FC of three independent biological replicates was used to calculate the average FC, the standard deviation (SD) and the standard error (SE), shown in the error bars.

### Nucleus-Cytoplasm fractionation

Nucleus-Cytoplasm fractionation was performed as described in Conrad & Ørom, 2017 to evaluate the subcellular localization of the circular and the linear isoforms transcribed from p-circ overexpression constructs. All centrifugation steps were carried out at 4°C and all buffers were ice cold.

Briefly, HEK 293T cells, previously transfected and induced as indicated above, were detached by adding 0.5 ml of 0.25 % Trypsin solution (Gibco) and incubating at 37°C for 5 min. The trypsinization reaction was then inactivated adding 1.5 ml of HEK maintaining media. Cell suspension was transferred into a 15 ml falcon tube, spun for 5 min at 200 × g in a tabletop centrifuge and the supernatant was aspirated. Cell pellet was resuspended in 10 ml PBS and spun at 200 × g for 5 min. The supernatant was removed, cell pellet was resuspended in 1 ml PBS and transferred to a 1.5 ml Eppendorf tube, spun at 200 × g in a microcentrifuge for 2 min and the supernatant was carefully removed again. 400 μl of Igepal lysis buffer (10 mM Tris pH 7.4, 150 mM NaCl, 0.15 % Igepal CA-630) was added to the pellet, gently pipetted up and down 3–5 times to resuspend the cells, and the solution was incubated on ice for 5 min. The cell lysate was gently transferred in a new Eppendorf tube and gently overlayed on top of 1 ml sucrose buffer (10 mM Tris pH 7.4, 150 mM NaCl, 24 % sucrose). The solution was centrifuged at 3500 × g for 10 min and the supernatant, containing the cytoplasmic fraction, was transferred in another tube and cleared again by centrifugation at 14,000 × g for 1 min. Then, 1/7 of the cytoplasmic fraction was resuspended in TRIZOL (Thermo Fisher). Instead, the pellet obtained after the centrifugation in sucrose buffer, corresponding to the nuclear fraction, was all resuspended in TRIZOL. RNA was extracted and, upon removal of residual genomic and plasmidic DNA, RNA concentration was quantified and iso-volumetric quantities (never exceeding 500 ng) from nuclei and cytoplasmic fractions were collected and retrotranscribed. For qPCR analysis, the logarithm to the base of 2 of 7 was subtracted to all the Ct means of the cytoplasmic fractions, to account for dilution coefficient of the cytoplasmic lysate compared to the nuclear one. Then, the 2^-Ct mean was calculated for nuclear and adjusted cytoplasmic fractions and their sum was used as reference to calculate the relative percentage of transcript in the specific compartment.

### Live imaging of Pepper-tagged RNAs

HEK 293T cells, previously transfected and induced as indicated above, were incubated for 5-30 minutes in FluoroBrite^TM^ DMEM medium (Gibco) supplemented with MgSO_4_ 5 mM and HBC620 0.5 μM (FR biotechnology^TM^) following manufacturer instructions. For experiments carried out in stress condition, 0.5 mM NaAsO_2_ (Sigma-Aldrich) was added to the imaging medium and incubated for 1 hour after HBC620 treatment to visualise RNAs and proteins during oxidative stress response. **RNase R treatment on live cells**

HEK 293T cells cultured and transfected in 8-well Nunc™ Lab-Tek™ II Chambered Coverglass as previously described, were first washed twice in PBS and then permeabilized in Triton X-100 0.05%/DPBS for 5 minutes (Ganassi et al., 2016). Subsequently, cells were washed again in PBS and incubated with imaging medium (MgSO_4_ 5 mM; HBC620 0.5mM in FluoroBrite^TM^ DMEM) for 15 minutes. Finally, cells were treated with 10 U of RNase R (Biosearch technologies, RNR07250) per well and imaged for 4 hours (Koppula et al., 2022) at a temporal resolution of 10 min/frame.

To evaluate RNase R activity, images were analysed using Fiji-ImageJ software: briefly, bleach correction with a Simple Ratio algorithm was applied on HBC620 signal and the polygon selection tool was used to generate ROIs and select HBC620 positive cells. The “Analyze > Measure” function was then used to calculate mean fluorescence intensity at time points corresponding to 0, 120, 180 and 240 minutes of RNase R treatment. The mean intensities of all the time points were then normalized on the value of the first time point. Samples were compared and tested for statistical significance with a two tails unpaired Student’s t-test.

For RNA quantification during RNase R treatment, cells were treated in the same conditions and collected in TRIZOL at each time point. RNA was extracted and DNase-treated as described above. 500 ng of RNA at time point 0, and an equal volume of RNA from the other time points for each condition were retrotranscribed with PrimeScript RT Master Mix (Takara Bio) to account for minimal cell death during RNase incubation. The qPCR results are expressed as 2^-dCt, were dCt is the difference between each time point Ct and the time point 0 Ct.

### Differentiation of mouse Embryonic Stem Cells (mESCs)-derived MNs

Mouse embryonic stem cells were cultured and differentiated into spinal MNs as described in Capauto et al, 2018; D’Ambra et al, 2024; Wichterle & Peljto, 2008. Briefly, cells were maintained in culture with mESC medium, (EmbryoMax DMEM; 15% Embryonic stem-cell FBS, ThermoFisher Scientific; 1% EmbryoMa× 100X nucleosides, Sigma-Aldrich; 1% EmbryoMax non-essential amino acids, Sigma-Aldrich; 2-mercaptoethanol for ES cells, Sigma-Aldrich; 2 mM L-glutamine, Sigma-Aldrich; 1% penicillin-streptomycin, Sigma-Aldrich) supplemented with ESGRO Recombinant Mouse LIF Protein (Chemicon), FGFR inhibitor PD173074 (Sigma-Aldrich P2499) and GSK-3 Inhibitor XVI (Sigma-Aldrich 361559).

For motoneuronal differentiation, embryoid bodies (EBs) were obtained by culturing mESCs in ADNFK medium (1:1) Advanced DMEM/F12 (Gibco):Neurobasal medium (Gibco), 10% Knock Out Serum Replacement (Gibco), 1% GlutaMAX, 1% 2-mercaptoethanol, 1% Pen/Strep. After two days, ADNFK medium was supplemented with 2% B27 Supplement (Gibco), 1μM RA (Sigma Aldrich) and 0.5 μM SAG (Merck Millipore). On day 5, ADNFK medium was supplemented with 2% B27 Supplement and 5 ng/mL GDNF (Peprotech). On day 6, EBs were dissociated: first, EBs were incubated with 20U/ml Papain (Worthington Biochemical Corporation), agitated for 5 minutes by hand, then blocked with 10 mg/ml ovomucoid inhibitor (Worthington Biochemical Corporation) for 5 min. Cells were then left to precipitate by gravity and supernatant was removed. Single cells were then dissociated by pipetting in PBS supplemented with 0.4% Glucose (Sigma), 2.5% horse serum (Thermofisher scientific), 2% B27, 3 mM MgCl_2_, Deoxyribonuclease I (Sigma-Aldrich, 25μg/ml) and plated on 0.01% poly-L-ornithine (Sigma-Aldrich), Laminin 20 μg/mL (Sigma) coated glass coverslips. MNs were maintained in culture with N2B27 medium (50 % DMEM/F-12 Ham, 50 % Neurobasal Medium, 1% GlutaMAX Supplement, 1% 2-mercaptoethanol, 1% non-essential amino acids, 0.5 % penicillin-streptomycin) supplemented with 2% N-2 supplement (Gibco), 1% B-27 supplement serum free (ThermoFisher Scientific), 200 ng/mL L-ascorbic acid (Sigma-Aldrich), 20 ng/mL BDNF (Peprotech), 10 ng/mL GDNF (Peprotech) and 10 ng/mL CNTF (Peprotech) and 10 mM ROCK inhibitor (Y-27632 dihydrochloride; Sigma-Aldrich).

For experiments carried out in stress condition, NaAsO_2_ at a final concentration of 0.5 mM was added to the neuronal medium and incubated for 1 hour.

### Differentiation of induced Pluripotent Stem cells (iPSCs)-derived MNs

Human iPSCs were maintained and differentiated in spinal MNs as described in De Santis et al, 2018. Briefly, iPSCs were dissociated to single cells with Accutase (Thermo Fisher Scientific) and plated in Nutristem-XF/FF medium (Biological Industries) supplemented with 10 μM ROCK inhibitor (Enzo Life Sciences) on Matrigel (BD Biosciences) at a density of 100′000 cells/cm^2^. The day after, differentiation was induced by adding 1 μg/ml doxycycline (Thermo Fisher Scientific) in Nutristem without bFGF and TGFβ (Biological Industries) in order to drive the expression of NIL (Ngn2-F2A-Isl1-T2A-Lhx3) cassette. After 48 hours of doxycycline induction, medium was changed to Neurobasal/B27 medium (Neurobasal Medium, Thermo Fisher Scientific, supplemented with 1X B27, Thermo Fisher Scientific, 1X Glutamax, Thermo Fisher Scientific, 1X NEAA, Thermo Fisher Scientific, and 0.5X Penicillin/Streptomycin, Sigma Aldrich), containing 5 μM DAPT and 4 μM SU5402 (both from Sigma Aldrich). At day 5, cells were dissociated with Accutase (Thermo Fisher Scientific) and plated on Matrigel (BD Biosciences) coated dishes. 10 μM ROCK inhibitor was added for the first 24 hours after dissociation. Neuronal cultures were maintained in neuronal medium (Neurobasal/B27 medium supplemented with 20 ng/ml BDNF, 10 ng/ml GDNF, both from PeproTech, and 200 ng/ml l-ascorbic acid, Sigma Aldrich).

For experiments carried out in stress condition, NaAsO_2_ at a final concentration of 0.5 mM was added to the neuronal medium and incubated for 1 hour.

### BaseScope^TM^ single molecule Fluorescent *in situ* Hybridization (smFISH) of circRNAs coupled with Immunofluorescence (IF)

mESCs derived MNs were plated on pre-coated 12 mm diameter coverslips and fixed in 4% paraformaldehyde (Electron Microscopy Sciences, Hatfield, PA) for 20 min at 4°C. Dehydration step with ice-cold Ethanol series (50%, 70%, 100%) was performed in order to store cells at -20°C in absolute ethanol until use. Detection of circ-Hdgfrp3 was performed via Basescope™ assay (Advanced Cell Diagnostics, Bio-Techne) as previously described in D’Ambra et al, 2024 with custom produced probes (Advanced Cell Diagnostics, ref. 703021) designed to specifically target its backsplicing junction. Briefly, fixed cells were permeabilized with Protease III (diluted 1:15; ref. 322381) before hybridization with the circ-Hdgfrp3-specific probes (Advanced Cell Diagnostics, Bio-Techne, cat#703021), at 40°C for 2 hours. Amplification and detection steps were performed following the manufacturer’s instructions using Basescope™ detection reagents V2– RED (ref. 323910). After each amplification step, three washes were performed with 300 ml of 1X RNAscope Wash Buffer Reagents (ref. 310091) for 5 minutes at room temperature.

FISH staining was combined with IF incubating the cells with the anti-DCP1A (ab183709) and anti-TIAR (BD Transduction MS BD 610352) primary antibodies used 1:100 and 1:200 in blocking solution 3% BSA and 0.2% Triton X-100 for 1 hour at room temperature. After three washes in PBS, cells were labelled with secondary antibodies: Goat anti-Rabbit 647 (ThermoFisher Scientific cat#A32795) diluted 1:300 in 1% goat serum/1% donkey serum/ PBS for 45 minutes at room temperature. Lastly, the nuclei were counterstained with DAPI solution (1 μg/mL/PBS; Sigma-Aldrich, D9542) for 5 minutes at room temperature and then the coverslips were mounted using ProLong Diamond Antifade Mountant (Thermo Fisher Scientific, P-36961).

### smFISH of long non-coding RNAs coupled with IF

iPSCs derived MNs were plated on 12 mm diameter coverslips coated with Geltrex (Thermofisher Scientific) and fixed in 4% paraformaldehyde (Electron Microscopy Sciences, Hatfield, PA) for 10 min at room temperature. Cells were stored at -20°C in absolute ethanol until used upon dehydration step with ice-cold Ethanol series (50%, 70%, 100%).

nHOTAIRM1 was detected via smFISH with a mix of 18 biotinylated probes (Sigma) as described in Santini et al, 2021 and Vautrot et al, 2015. Briefly, cells were rehydrated by descendent ice-cold ethanol series (100%, 70%, 50%) and permeabilized in 0.05% Triton X-100/2 mM VRC (Sigma-Aldrich, R3380)/DPBS for 5 min. DPBS was replaced with 2X SSC buffer (300 mM NaCl; 30 mM sodium citrate in nuclease free water) and cells were incubated for 5 min. Finally, samples were incubated in pre-hybridization buffer (10% deionized formamide, Sigma-Aldrich, 47671; 2X SSC in nuclease free water) for 15 min at 37°C, followed by an over night incubation at 37°C in a slide hybridizer machine (ACD HybEZ™ II Hybridization System) with hybridization buffer (10% deionized formamide; 2X SSC; 10% w/v Dextran sulfate, Sigma-Aldrich, D8906, 2 mM vanadyl ribonucleoside complexes (VRC), Sigma-Aldrich, R3380, in nuclease free water) completed with the biotinylated probes at a final concentration of 50 nM each. The next day, cells were washed twice with 2X SSC for 5 min first at 37°C and then at RT. SSC buffer was then replaced with TN buffer (Tris HCl pH 7.5 10 mM; NaCl 5 mM in nuclease free water), incubated at RT for 10 min. Lastly, biotinylated oligoes were stained with 1:200 diluted Alexa Fluor^TM^ 568-conjugated streptavidin (Invitrogen™ S11226) incubated in 4% w/v BSA (Sigma-Aldrich, A2153)/TN buffer for 1-2 hours at room temperature in a humid box.

When FISH was combined with IF, cells were washed twice with TN buffer and once with DPBS and then were incubated with primary antibodies (anti-FUS, Bioss Antibodies bs-2980R, anti-FUS, Santa Cruz sc-47711, anti-G3BP1, Sigma GW22382A, anti-G3BP1, Sigma PLA0231, anti-TIAR, BD Transduction MS BD 610352, anti-DCP1A, ab183709) diluted in 1% w/v BSA/DPBS for 1 hour at room temperature. Then, upon three washes in DPBS, cells were incubated with 1:300 diluted secondary antibodies (Goat anti-Mouse Alexa Fluor^TM^ 488, Invitrogen A-11029; Goat anti-rabbit Alexa Fluor^TM^ 488, Invitrogen A-11008; Goat anti-Chicken Alexa Fluor^TM^ Plus 488, Invitrogen A32931, Donkey anti-mouse Alexa Fluor^TM^ 647, Invitrogen A-31571) in 1% w/v BSA/DPBS for 45 min at room temperature. Cells were then washed three times with DPBS, nuclei were counterstained with DAPI solution (Sigma, D9542; 1μg/ml/PBS) for 5 min at room temperature and coverslips were mounted with Prolong Diamond Mounting Media (ThermoFischer Scientific, P-36961).

### Image acquisition

Fixed mESC-derived MNs were imaged with confocal Olympus IX73 microscope equipped with a Crestoptics X-LIGHT V3 spinning disk system and a Prime BSI Express Scientific CMOS camera using UPlanSApo 100x (NA 1.45) oil objective and were collected with the MetaMorph software (Molecular Devices). The Z-stack confocal microscopy images were taken automatically (200 nm Z-spacing).

Fixed iPSCs-derived MNs were imaged on a Nikon Instrument A1 Confocal Laser Microscope equipped with a 1.49 NA 100x objective (Plan Apo VC 100x Oil DIC N2, Nikon, Tokyo, Japan). Confocal images were collected with NIS-Elements AR software (Nikon): ND Acquisition module was used for multipoint acquisition of Z-stack images (150-175nm Z-spacing) of 4 μm thickness.). Confocal images were collected with NIS-Elements AR software (Nikon): ND Acquisition module was used for multipoint acquisition of Z-stack images (150-175nm Z-spacing) of 4 μm thickness. All live imaging experiments were performed using an Eclipse Ti2-E Inverted Microscope equipped with the Nikon Super Resolution System (N-STORM & N-SIM), with a 1.49 NA 100x objective (Apo TIRF 100x Oil, Nikon, Tokyo, Japan) and with a 3D EX V-R 100x/1.49 Grating Block. Cells were kept at 37°C and with 5% CO_2_ supply with a live imaging control system (Tokai Hit, INU) for all the experiment duration. Widefield and SIM images were collected with NIS-Elements AR software (Nikon): ND acquisition module was used for multipoint and time-lapse images collection. Specifically, time-lapses with duration of 10-15 min and “no delay” (∼11 sec) interval were collected for higher temporal resolution acquisitions and time-lapses with duration of 1 hour and 1 to 5 minutes interval, depending on the number of selected multi-points, were collected for acquisitions upon NaAsO_2_ treatment. To reconstruct SIM acquisitions, the three reconstruction parameters illumination modulation contrast, high-resolution noise suppression and out of focus blur suppression were adopted to generate consistent Fourier transform. Images with a reconstruction score of 8 were selected for sub-structural analysis.

Occasionally, the Denoise.ai and the Clarify.ai deconvolution algorithms available on NIS-Elements AR software (Nikon), were used to post-process Widefield time-lapse acquisitions for representative images.

### Co-localization analysis and particles measurements on fixed samples

Fiji-ImageJ open source software was used for analysis on confocal images of immunofluorescence and FISH experiments.

For DCP1A/circ-Hdgfrp3 co-localization in mESC-derived MNs all images were analysed as it was previously described in D’Ambra et al, 2021. Briefly, Z-stacks were processed with Laplacian of Gaussian filter (Ballarino et al., 2018; Sage et al., 2005) and intensity threshold, contrast and brightness were adjusted using the ImageJ software. The co-planarity evaluation of the signals was performed combining the fluorescence distributions of each channel, recorded in the main greyscale value (expressed as arbitrary units) along Z-planes obtained from the ImageJ Plot Z-axis profile plugin and the count of co-localizing signal were performed manually.

For object-based co-localization analysis between the lncRNA nHOTAIRM1, G3BP1 and FUS^P525L^, and between endogenous DCP1A and exogenous GFP-DCP1A, Moment’s algorithm was exploited (Tsai, 1985) to create binary masks for FISH and IF channels. Otsu thresholding algorithm (Otsu, 1979) was used to create nuclei binary masks starting from DAPI signal. The Image Calculator command was then used to subtract nuclei masks to nHOTAIRM1, G3BP1, FUS^P525L^ and DCP1A channels in order to account only for cytoplasmic signal. To detect co-localizing particles in the cytoplasm, the Image Calculator “AND” function was then applied to create secondary masks resulting from the intersection of the pixels between: nHOTAIRM1 and G3BP1; nHOTAIRM1 and FUS^P525L^; nHOTAIRM1, G3BP1 and FUS^P525L^; FUS^P525L^ and G3BP1 or between endogenous DCP1A and exogenous GFP-DCP1A. “Analyze particles” function was used to count number of total particles and number of particles in the intersection masks, thus percentages of co-localization were obtained as a ratio between particles in the intersection mask/total number of particles.

For pixel-wise co-localization between nHOTAIRM1 and DCP1A, nHOTAIRM1 and TIAR and TIAR and DCP1A, a Median filter radius = 2 was applied and the Plug-in JACoP of Fiji-Imagej was used to calculate Pearson’s co-localization coefficients (PCC) between the corresponding channels (Bolte and Cordelieres, 2006). A PCC ranging from -1 to 0 was considered as “anti-correlation”; a PPC ranging from 0 to 0,1 was described as “poor correlation”; a PPC > 0,1was described as “correlation”

### Single-particle tracking and sub-structural distribution of Pepper-tagged RNAs and GFP-tagged proteins in live cells

TrackMate plug-in on Fiji-ImageJ was used for single-particle tracking of Pepper-tagged RNAs interacting with G3BP1-, FUS^P525L^- or GFP-DCP1A (Ershov et al., 2022; Tinevez et al., 2017). Representative ROIs were selected from time-lapse acquisition with duration of 10-15 min and “no delay” (∼11 sec) interval, collected with widefield microscopy. DoG Detector algorithm was selected and an “estimated object diameter” of 1 μm was chosen to filter for the objects to track, while “Quality threshold” was determined case by case. Linear Assignment Problem (LAP) tracking algorithm (Jaqaman et al., 2008) was then exploited with the following configuration options:

- Frame to frame linking: 1 μm;
- Gap-closing max distance: 1 μm;
- Max frame gap: 3;
- Max distance for track segment splitting: 1 μm;
- Max distance for track segment merging: 1 μm;

Finally, “track tables” recording single spots IDs and xy-coordinates frame-by-frame were extrapolated and used to generate 3D scatter plots with the Origin Lab software. In particular, x-axis was assigned to x-coordinates, y-axis was assigned to time position and z-axis was assigned to y-coordinates.

To determine the sub-structural distribution of circ-HDGFRP3 in FUS^P525L^ aggregates, profiles of the signal distribution were generated along an arbitrary line with the function “Analyze > Plot Profile” of the Fiji-ImageJ menu. The axial resolution of representative SIM images was estimated thanks to the Image Decorrelation Analysis plugin on Fiji-ImageJ (Descloux et al., 2019).

### Pixel-wise and Object-based co-localization analysis in live samples

To evaluate the association throughout time between the RNAs and the GFP-tagged proteins upon oxidative stress induction a pixel-wise co-localization approach was chosen. A custom ImageJ macro was used to calculate Pearson’s and Manders’ coefficient between the RNA (HBC620) and the protein (eGFP) signals frame-by-frame from time-lapses of 1 hour duration and 1-5 min interval. Briefly, bleach correction with a Simple Ratio algorithm was applied on both HBC620 and GFP channels, then a DoG filter was applied to both channels generating a noise reduced image (Gaussian blur σ = 2) and a background image (Gaussian blur σ = 10) and subtracting the two with the Image Calculator “Subtract” function. A ROI was then selected to analyse only cells in the field of view double positive for HBC620 and eGFP signals and the Plug-in JACoP of Fiji-ImageJ was used to calculate co-localization coefficients on each frame (Bolte and Cordelieres, 2006). To account for the increase in fluorescence intensity as the G3BP1 and FUS^P525L^ proteins condensate upon stress, thresholder Manders’ Coefficient was evaluated when analysing nHOTAIRM1/G3BP1, nHOTAIRM1/FUS^P525L^, ROMO1/G3BP1 or ROMO1/FUS^P525L^ time-lapses (Manders et al., 1993). Threshold values for both channels were determined for each acquisition depending on the fluorescence intensity.

Conversely, to analyse the association between FUS^P525L^ aggregates and Pepper-tagged RNAs, an object-based approach was chosen. Briefly, bleach correction with a Simple Ratio algorithm was applied on both HBC620 and GFP channels, then a LoG filter was applied to both channels taking advantage of the LoG 3D plug-in on ImageJ (Ballarino et al., 2018; Sage et al., 2005). Intermodes thresholding algorithm (Prewitt and Mendelsohn, 2006) was then exploited to create binary masks of

HBC620 and GFP channels and the Image Calculator “AND” function was applied to create a secondary masks resulting from the intersection of the pixels between the two masks. “Analyze particles” function was used to count number of total RNA and FUS^P525L^ particles and number of particles in the intersection masks in each time frame, thus fraction of co-localizing particles was obtained as a ratio between number of particles in the intersection mask/total number of RNA (HBC620) particles per time interval.

## DATA AVAILABILITY

This study has not generated data that requires deposition in a public database.

## AUTHOR CONTRIBUTION

I.B. conceptualized the project. I.B., D.M., E.C. and P.L. supervised and coordinated the experiments. IB., G.R and P.L provided founding. E.V. and F.C. performed live-imaging experiments. E.V., F.C., L.S.M. D.M. and P.T. performed cloning and molecular biology assays. F.C. and P.T. cultured and differentiated iPSCs. E.D.A. cultured and differentiated mESCs. E.V. and E.D.A. performed and analysed FISH and IF experiments. E.V. analysed live imaging experiments. I.B. and E.V. wrote the original manuscript with contribution of all the authors.

## ACKNOWLEDGMENTS

We thank Prof. Alberto Diaspro and Dr. Paolo Bianchini (Nanoscopy & NIC@IIT, Istituto Italiano di Tecnologia), Dr. Michele Oneto (Nikon Imaging Center) and Marco Scotto (Molecular Microscopy and Spectroscopy, Istituto Italiano di Tecnologia) for experimental support in confocal and SIM microscopy. We are also grateful to Dr. Giuseppe Vicidomini, Dr. Eleonora Perego and Sabrina Zappone (Molecular Microscopy and Spectroscopy, Istituto Italiano di Tecnologia) for useful discussion and suggestions. The authors would also like to thank the “RNA Technologies” Flagship at IIT.

This work was supported by grants from European Research Council [ERC-2019-SyG 855923-ASTRA] and from Italian Association for Cancer Research [AIRC IG 2019, Id. 23053] to I.B.; from PRIN 2017 [id. 2017P352Z4] and from EU within the MUR PNRR “National Center for Gene Therapy and Drugs based on RNA Technology” [project no. CN00000041 CN3, Spoke #3 “Neurodegeneration” and Spoke #6 “RNA Drug Development”] to I.B. and P.L.; from PRIN 2022 [id. 2022BYB33L] and from Consiglio Nazionale delle Ricerche-CNR [projects DBA.AD005.225-NUTRAGE-FOE2021 and DSB.AD006.371-InvAt-FOE2022] to P.L.; and from European Innovation Council (EIC) through its Pathfinder Open Programme, project ivBM-4PAP [id. 101098989], together with intramural IIT fundings to G.R.

